# Co-Phosphorylation Networks Reveal Subtype-Specific Signaling Modules in Breast Cancer

**DOI:** 10.1101/2020.05.04.073148

**Authors:** Marzieh Ayati, Mark R Chance, Mehmet Koyutürk

**Affiliations:** Department of Computer Science, University of Texas Rio Grande Valley, Edinburg, TX, 78531, USA; Department of Nutrition, Case Western Reserve University, Cleveland, OH; Center for Proteomics and Bioinformatics, Case Western Reserve University, Cleveland, OH; Case Comprehensive Cancer Center, Case Western Reserve University, Cleveland, OH; Department of Computer and Data Sciences, Case Western Reserve University, Cleveland, OH, 44106, USA

## Abstract

**Motivation:** Protein phosphorylation is a ubiquitous mechanism of post-translational modification that plays a central role in cellular signaling. Phosphorylation is particularly important in the context of cancer, as down-regulation of tumor suppressors and up-regulation of oncogenes by the dysregulation of associated kinase and phosphatase networks are shown to have key roles in tumor growth and progression. Despite recent advances that enable large-scale monitoring of protein phosphorylation, these data are not fully incorporated into such computational tasks as phenotyping and subtyping of cancers.

**Results:** We develop a network-based algorithm, CoPPNet, to enable unsupervised subtyping of cancers using phosphorylation data. For this purpose, we integrate prior knowledge on evolutionary, structural, and functional association of phosphosites, kinase-substrate associations, and protein-protein interactions with the correlation of phosphorylation of phosphosites across different tumor samples (a.k.a co-phosphorylation) to construct a context-specific weighted network of phosphosites. We then mine these networks to identify subnetworks with correlated phosphorylation patterns. We apply the proposed framework to two mass-spectrometry based phosphorylation datasets for breast cancer, and observe that (i) the phosphorylation pattern of the identified subnetworks are highly correlated with clinically identified subtypes, and (ii) the identified subnetworks are highly reproducible across datasets that are derived from different studies. Our results show that integration of quantitative phosphorylation data with network frameworks can provide mechanistic insights into the differences between the signaling mechanisms that drive breast cancer subtypes. Furthermore, the reproducibility of the identified subnetworks suggests that phosphorylation can provide robust classification of disease response and markers.

**Availability and implementation:** CoPPNet is available at http://compbio.case.edu/coppnet/

## 1 Introduction

Protein phosphorylation is a ubiquitous mechanism of post-translational modification observed across cell types and species, and plays a central role in cellular signaling. Phosphorylation is regulated by networks composed of kinases, phosphatases, and their substrates. Phosphorylation is particularly important in the context of cancer, as down-regulation of tumor suppressors and up-regulation of oncogenes (often kinases themselves) by dys-regulation of the associated kinase and phosphatase networks are shown to have key roles in tumor growth and progression [1, 2]. To this end, characterization of signaling networks enables exploration of the interconnected targets leading to the development of kinase inhibitors to treat a variety of cancers [3, 4]. In response to the growing need for large-scale monitoring of phosphorylation, advanced mass spectrometry (MS)-based phospho-proteomics technologies have exploded. These technologies enable simultaneous identification and quantification of thousands of phosphopeptides and phosphosites from a given sample [5].

MS-based phospho-proteomics screens create a great opportunity to discover biology that may not be observed in transcriptomic and proteomic data [6]. Indeed, recent research shows that, as compared to gene expression, data on post-transcriptional modifications can be more useful in subtyping cancers. As a striking example, monitoring of the specific phosphorylation pathways reveals a novel breast cancer subtype that is unique to the phospho-proteomics and cannot be captured based on DNA mutations, mRNA-level expression, or protein expression [7].

Although phospho-proteomics provides a critical data source to model signaling pathways, systematic methods for network analysis of phospho-proteins and phosphosites are relatively scarce. Since most of the methods designed for genomics and general proteomics are not designed to handle the complexity of phospho-proteomics, phospho-proteomic analyses are often centralized at the protein level. However, due to the *many-to-one* mapping from phosphosites to proteins (i.e. each protein may have multiple phosphorylation sites), and also multi-layer annotations (e.g. regulatory function of phosphosites and kinase-phosphosite associations), novel approaches are needed to fully leverage the richness of the data. To enable analysis of phospho-proteomic data at the level of phosphorylation sites and the relationships between these sites, we propose CoPPNet, a network-based algorithm for the analysis of phosphoproteomic data, which offers the following innovations: (i) Construction of a PhosphoSite Functional Association (PSFA) network that represents the functional relationship among individual phosphosites. In order to create PSFA network, we incorporate known structural, evolutionary, and functional associations between phosphosites, protein-protein interactions, and kinase-substrate associations. (ii) Utilization of the PFSA network in the identification of phosphorylation modules in breast cancer, through filtering of phosphosite pairs that are potentially functionally associated. CoPPNet accomplishes this by assigning co-phosphorylation (Co-P) based weights to the edges in PFSA network, where Co-P quantifies the similarity of the phosphorylation patterns of phosphosites across different breast cancer samples. We have recently introduced the notion of co-phosphorylation and used it in the context of predicting kinase-substrate associations, showing that it significantly enhances the coverage and accuracy of prediction methods over those that utilize static data such as sequences, structures, and generic networks [8]. Conceptually, Co-P is similar to gene co-expression, which has been shown to be effective in many biomedical applications [9, 10]. (iii) Development of a scoring scheme accompanied by an algorithm to identify co-phosphorylated signaling modules from this weighted PSFA network.

We test the proposed framework in the context of *unsupervised* identification of subtype-specific signaling modules in breast cancer. For this purpose, we apply CoPPNet on two independent public phospho-proteomics datasets for breast cancer (BC). Breast cancer is categorized into 4 molecular subtypes: Luminal A, Luminal B, HER2-enriched and triple-negative (Basal-like). Among the subtypes, Luminal A has the greatest survival, and Basal has the poorest survival [11]. While constructing the weighted PSFA network and identifying co-phosphorylation modules on this network, we do not use any information on the samples’ clinically determined subtypes.

Our results show that the statistically significant modules identified by CoPPNet are reproducible between the two independent datasets and can capture the differential phosphorylation between breast cancer subtypes. The identified subtype-specific signaling modules have the potential to provide significant insights into the disruption of signaling processes in different cancer subtypes, and can be employed in developing subtype specific therapeutic targeting strategies for breast cancer.

## 2 MATERIALS AND METHODS

The workflow of the proposed framework for unsupervised identification of co-phosphorylation (Co-P) modules is shown in Figure 1. As seen in the figure, we first construct a network to model the functional relationship between phosphorylation sites. For this purpose, we incorporate available knowledge on functional associations between phosphosites, kinase-substrate associations and protein-protein interactions, and integrate these knowledge into a PhosphoSite Functional Association (PSFA) network. Subsequently, we utilize a module identification algorithm to identify sub-networks of the PSFA network that are composed of highly co-phosphorylated phosphosites (called *Co-P modules*). The premise of this approach is that, pairs of phosphosites whose phosphorylation is related to a specific cancer subtype will exhibit co-variation across different samples. For this reason, we expect that Co-P can highlight subtype-specific signaling modules even if subtype information is not available for the samples that are used to compute Co-P.

**Figure 1:**
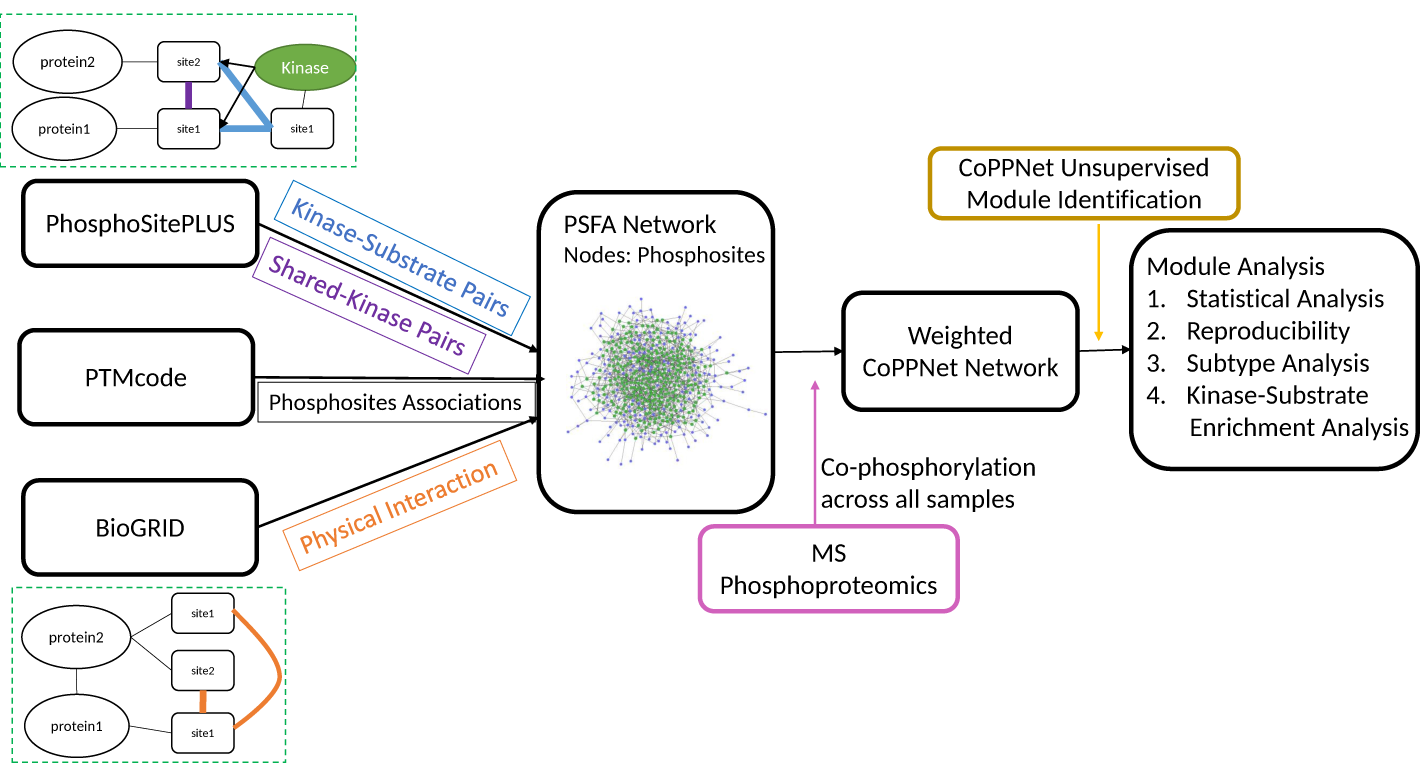
Workflow of CoPPNet. We first construct a PSFA network to represents the functional relationship among phosphosites, by utilizing generic kinase-substrate association, phosphosites associations and protein-protein interaction data. The nodes of the PSFA network represent phosphosites and the edges represent (1) kinase-substrate association, 2) phosphosites targeted by a common kinase, (3) functional associations between phosphosites, (4) physical interaction between proteins harboring the sites. For a given phosphorylation dataset collected from multiple cancer samples, we weigh the edges of the PSFA network based on the co-phosphorylation (Co-P) of pairs of sites across these samples. Then, we identify Co-P modules as sub-networks composed of heavy edges in this weighted network. Finally, we comprehensively assess the significance, reproducibility, subtype-specificity, and biological relevance of the Co-P modules.

To assess the biological significance of the identified significant modules, we comprehensively evaluate their statistical significance and investigate the reproducibility of significant modules by utilizing a dataset that comes from a different patient cohort. Subsequently, we assess the differential phosphorylation of the sites in the signaling modules between different subtypes and perform pathway enrichment analysis and kinase enrichment analysis on these modules to annotate the modules.

### PhosphoSite Functional Association (PSFA) Network

We define a PhosphoSite Functional Association (PSFA) network as a network that represents *potential* functional relationships between pairs of phosphosites. This network serves the purpose of filtering out the search space for pairs of phosphosites whose co-phosphorylation may reveal their functional relationship in the context of a specific process (e.g., dysregulation of a signaling pathway in the progression of a certain cancer subtype). In PSFA network *G*(*V, E*), *V* denotes the set of nodes in the network, each of which represents a phosphosite; thus a protein is represented by multiple nodes in the PSFA network. The edge set *E* denotes the set of pairwise functional relationships between phosphosites, where an edge *s*_*i*_*s*_*j*_ ∈ *E* between phosphosites *s*_*i*_, *s*_*j*_ ∈ *V* may represent one of the following relationships:

- **Functional, Evolutionary, and Structural Association between Phosphosites (FES).** PTMCode is a database of known and predicted functional associations between phosphorylation and other post-translational modification sites [12]. The associations included in PTMCode are curated from the literature, inferred from residue co-evolution, or are based on the structural distances between phosphosites. We utilize PTMcode as a direct source of functional, evolutionary, and structural associations between phosphorylation sites.
- **Kinase-Substrate Association (KSA).** If phosphosite *s*_*i*_ is a target of kinase *p*_*k*_ and *s*_*j*_ is a phosphosite on kinase *p*_*k*_, then there is an edge between *s*_*i*_ and *s*_*j*_ in the PFSA network. We call these edges *KSA edges*. This relationship indicates potential functional association between *s*_*i*_ and *s*_*j*_ since the regulation of kinase *p*_*k*_ through phosphorylation of *s*_*j*_ may influence *p*_*k*_’s action on *s*_*i*_. In our experiments, we use PhosphositePLUS as the main source of information for kinase-substrate association [13].
- **Phosphosites Targeted by Common Kinase (TCK).** If phosphosites *s*_*i*_ and *s*_*j*_ (which may be on the same protein or on different proteins) are targeted by kinase *p*_*k*_, then we call them a *shared-kinase pair* and include an edge between *s*_*i*_ and *s*_*j*_ in the PSFA network. We call these edges TCK edges. We include TCK edges in the PSFA network since the activity of *p*_*k*_ in a specific process may influence the phosphorylation of both *s*_*i*_ and *s*_*j*_, which may be captured by their co-phosphorylation. Indeed, studies have shown that the substrates of a protein kinase can have significant similarity in terms of their biological functions [14].
- **Protein-Protein Interaction (PPI)**. If two proteins *p*_*f*_ and *p*_*r*_ physically interact, for any site *s*_*i*_ is on *p*_*f*_, and site *s*_*j*_ is on protein *p*_*r*_, then there is an edge between *s*_*i*_ and *s*_*j*_ in the PSFA network. We call these edges *PPI edges*. We include PPI edges in the PSFA network, since these edges may capture functional relationships and post-transcriptional modifications beyond phosphorylation, and may remedy the sparse and incomplete nature of existing kinase-substrate annotations. In our experiments, we use the PPIs that are annotated as “physical” in the BIOGRID PPI database [15] to infer the PPI edges in the PFSA network.

The PSFA network is a generic network of potential functional associations between pairs of phosphosites. In the next section, we discuss how to assign weights to the edges of the PSFA network to represent the co-phosphorylation of pairs of phosphosites in a specific context.

### Assessment of Co-Phosphorylation

As with gene co-expression, correlated phosphorylation of phosphosites on proteins may be indicative of their functional relationship in a specific biological context [8]. Based on this premise, we use context-specific phosphorylation data, obtained from mass spectrometry based phospho-proteomics assays, to assess the co-phosphorylation (Co-P) of all pairs of phosphosites that are connected in the PSFA network. In gene co-expression analysis, Pearson’s correlation and mutual information are commonly used to assess linear and non-linear relations between the expression profiles of genes [16, 17]. Recognizing the benefits and shortcomings of each method, Song et al. [18] developed bi-weight mid-correlation as an alternative, and showed that it outperforms mutual information in terms of capturing biologically relevant relationships between genes. while being more robust to outliers than Pearson’s correlation. Motivated by these results, we use bi-weight mid-correlation to assess the Co-P of pairs of phosphosites.

### Identification of Co-Phosphorylation Modules

Given a weighted PSFA network *G*(*V, E, w*) associated with a specific phosho-proteomic dataset, our objective is to identify sub-networks of the PSFA network that are enriched in highly co-phosphorylated (positively or negatively) pairs of phosphosites. This problem is similar to the well-studied problem of identifying altered sub-networks, in which the nodes are scored based on their dysregulation (e.g., z-score indicating differential gene expression) in a given condition [19] or association with a disease (e.g., − log of the *p*-value of association) [20]. In this network, one or more connected sub-networks composed of high-scoring nodes are sought. In contrast, in our problem, scores are associated with edges, thus the problem is also similar to the infamous community detection problem in network analysis.

As with the altered sub-network identification problem, the key component of a solution to the problem is the definition of an objective function for scoring a given sub-network. Inspired by Newman’s definition of network modularity [21] and our adaptation of this measure to the identification of disease-associated modules [20], we here propose a modularity-based approach to scoring co-phosphorylation modules. In this approach, subnetworks are scored based on the difference between their total edge weight and their expected total edge weight under a reference model that takes into account the degree distribution of the network (in our case, the distribution of Co-P across the network). Namely, for a given set of phosphosites *Q* ⊆ *V*, we define the Co-P score of *Q* according to ℳ as

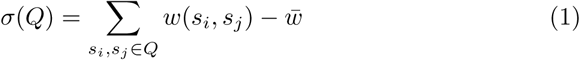

where 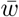 is the mean of the absolute values of Co-P across all pairs of phosphosites, and *w*(*s*_*i*_, *s*_*j*_) is Co-P of *s*_*i*_ and *s*_*j*_ if there is an edge between them and 0 otherwise. Thus, it penalizes the non-existent edges.

Having defined the Co-P score of a subnetwork as in Equation 1, given weighted PSFA network *G*(*V, E, w*), we search for subnetworks of *G* that maximize *σ*(*Q*). Since the *maximum-weight induced subgraph problem* is NP-hard [22], we use a greedy algorithm for this purpose. Namely, we search the network by starting from the phosphosite with the largest fold change, repeatedly examining the phosphosites in the neighborhood of the phosphosites so far in the subnetwork, and adding to the subnetwork the phosphosites that provide the best improvement of the subnetwork score. Once we identify a subnetwork with locally maximal Co-P score, we remove this subnetwork from *G* and use the greedy algorithm again to identify the next subnetwork with locally maximal Co-P score. We repeat this procedure until the entire network is exhausted, and sort all of the identified subnetworks (called Co-P modules) in decreasing order of their Co-P score. The pseudocode of the algorithm is provided in Supplementary material. We also compare the performance of this algorithm against two other state-of-the-art module identification algorithms: Girvan and Newman’s algorithm for the identification of communities [23] and the WGCNA algorithm for clustering gene co-expression networks [24, 25]. We observe that the subnetworks identified by other algorithms are less parsimonious and tend to be composed of sites that are on the same protein. CoPPNet identifies more parsimonious and statistically significant subnetworks by including a penalty term for non-extant edges in its objective function. Since the PFSA network does not include edges between sites on the same protein unless they are functionally associated, CoPPNet is able to identify signaling modules that span across multiple proteins. We report these results in detail in supplementary material.

### Assessment of Statistical Significance

To assess the statistical significance of all identified Co-P modules, we use two types of permutation tests. For this purpose, we use two null models: (i) randomize the weights of the edges of the PSFA network while preserving the topology of the network (thereby preserving the degree distribution of the phosphosites) to generate *N* permuted networks, and (ii) we permute the interactions while preserving the degree of phosphosites (we use *N* = 100 in the experimental results reported in the next section). On each of the permuted networks, we identify and rank Co-P modules using the algorithm described in the previous section. We then assess the statistical significance of each module identified on the original network by comparing its score against the scores of the subnetworks that are ranked at least as high as itself on the permuted networks. We also visualize the scores of the identified modules in the context of these cumulative empirical distributions. We pick the modules that are statisitically significant in terms of both null models for further analysis.

### Assessment of Subtype Specificity

Although the weights of edges in the PSFA network are computed using co-phosphorylation (Co-P), which is agnostic to the subtypes of the samples, Co-P captures the co-variation of phosphorylation levels of phosphosites across different samples. Therefore, the identified modules have the potential to be associated with subtype-relevant mechanisms. Motivated by this insight, we investigate if the identified Co-P modules are composed of phosphosites that exhibit differential phosphorylation between cancer subtypes. For this purpose, we assess the differential phosphorylation of each phosphosite in a module between different subtypes. We use standard *t*-tests to compare the distribution of relative phosphorylation level (with respect to the common reference) in different subtypes.

### Assessment of Predictive Ability

To assess the utility of identified modules in predicting subtypes, we train a support vector machine (SVM) based classifier on one dataset using the sites in the significant modules as features and assess the performance of this classifier in predicting subtypes on the other dataset. We compare the performance of these module-based features against a full model (incorporating all sites) and a model that incorporates all sites that are significantly deferentially phosphorylated (p¡0.05) between subtypes on the training dataset.

### Assessment of Reproducibility

We assess the *reproducibility of identified Co-P modules* by investigating the overlap between significant modules identified on independent datasets. To assess the overlap between two Co-P modules that are identified in two independent datasets, we use standard hypergeometric test. We assess the *reproducibility of subtype specificity* by computing the correlation between the fold changes of sites in the modules with respect to subtypes across the two datasets. We assess the significance of this correlation empirically using a permutation test.

### Kinase Substrate Enrichment Analysis

Kinase Substrate Enrichment Analysis (KSEA) seeks to identify kinases whose targets exhibit significantly altered phosphorylation levels in a given condition. KSEA scores each kinase based on the relative phosphorylation and dephosphorylation of its substrates (i.e fold change). In order to assess the value added by Co-P modules, we perform kinase enrichment analysis by restricting KSEA to the substrates that are in the significant modules as opposed to all phosphosites that are identified in the study. To infer the differential activity of kinases between subtypes, we compare the score of kinases which are computed using the fold change of target phosphosites across samples in different subtypes. We identify the kinases that are predicted to have different activity by KSEA using all sites vs. module-restricted sites and investigate the association of these kinases with survival using integrated gene expression data and survival information of 1809 patients from the Gene Expression Omnibus (GEO) [26].

### Protein Expression Analysis

We also investigate if protein phosphorylation data provide information on cancer substypes beyond what can be captured by protein expression. For this purpose, we utilize mass-spectrometry based protein expression data that is obtained from the samples that are used to obtain the phospho-proteomic data used in our computational experiments.

We utilize protein expression data in the following way: Using the phosphoproteomic data, we identify phosphosites in Co-P modules that are significantly differentially expressed (*p <* 0.05) between different subtypes. Subsequently, using proteomic data, we assess the differential expression of the proteins that harbor these significant phosphosites between different subtypes. If the protein that harbor the site is not identified in the protein expression data, we exclude them from the analysis.The result of this analysis is presented in Supplementary material.

## 3 Results and Discussion

### Datasets

#### Phosphoproteomics Data

We use two independent public quantitative mass spectrometry (MS) based phospho-proteomics datasets obtained from breast cancer (BC) Patient-Derived Xenografts (PDX).

- **Huang** *et al.* **data:** Huang *et al.* [27] used isobaric tags for relative and absolute quantification (iTRAQ) to identify 56874 phosphosites in 24 breast cancer PDX models. The clinically determined subtypes for the samples in this dataset are *Basal* for 10 samples, *Luminal* for 9 samples and *HER2-enriched* for 5 samples. We remove phosphosites with missing intensity values in any sample. This results in intensity data for 15780 phosphosites from 4539 proteins, where 13840 serines, 2280 threonines and 67 tyrosines are phosphorylated. Protein expression data for all of these samples is also available.
- **Mertin** *et al.* **data:** The NCI Clinical Proteomic Tumor Analysis Consortium (CPTAC) conducted an extensive MS based phospho-proteomics of TCGA breast cancer samples [7]. After selecting the subset of samples that have the highest coverage and filtering the phosphosites with missing intensity values in those tumors, the remaining data contained intensity values for 11018 phosphosites mapping to 8304 phosphoproteins in 20 tumors. This dataset contains 4 *Basal*, 9 *Luminal* and 7 *HER2-enriched* samples.

#### Functional, Evolutionary, and Structural Association between Phosphosites (FES)

We use PTMcode, a database for functional associations of post-translational modifications within and between proteins [12]. The functional association between PTM sites have been reported based on the literature survey, co-evolution of sites, structural proximity and if PTMs at the same residue and location are within PTM highly enriched protein regions. For our analysis, we just focus on the functional associations between phosphorylation sites of different proteins.

#### Kinase-Substrate Associations (KSAs)

We use PhosphoSitePLUS as a reference dataset for kinase-substrate associations [13]. PhosphoSitePLUS reported 9699 kinase-substrate association over 347 kinases.

#### Protein-Protein Interaction (PPI) Data

We use a generic human PPI network downloaded from BioGRID database at *https://thebiogrid.org* [15]. This network contains 194639 interactions among 18719 proteins.

The number of sites and edges in the final PSFA network and their types are shown in Table 1. This result suggests that all different types of edges contribute to the functional relevance of the phosphosites in the modules. Although there are more PPI edges in the PSFA network, TCK edges play an important role in the identification of signaling modules, since these edges induce cliques in the PFSA network. In this respect, CoPPNet implicitly identifies kinases whose targets exhibit enriched differential phosphorylation in specific subtypes. We elaborate on this feature of CoPPNet in the context of kinase enrichment analysis later in this section. The overlap between different types of edges is presented in Table S1.

**Table 1:** Number of phosphosites and edges in PSFA network and statistically significant modules

| Type of Edges<br># sites/ # edges | PSFA Network<br>9652/173772 | Module 1<br>91/4095 | Module 2<br>68/2026 |
| --- | --- | --- | --- |
| FES | 7999 | 1 | 6 |
| KSA | 3024 | 93 | 306 |
| TCK | 34857 | 4095 | 1714 |
| PPI | 133536 | 46 | 17 |

### CoPPNet identifies co-phosphorylation (Co-P) modules that are statistically significant and reproducible

We identify co-phosphorylated sub-networks on each of the two datasets using CoPPNet. We investigate the statistical significance of these subnetworks and visualize the results of this analysis in Figure 2(a). As seen in the figure, the two top-scoring subnetworks identified on both datasets have scores at least two standard deviation above the mean of the top subnetworks identified on 100 randomized networks. At a *q*-value threshold of 0.01, two of these subnetworks are detected to be statistically significant for each dataset. Note that since module identification is exhaustive, we don’t expect all the identified modules to be significant. In contrast, we observe that, with the exception of the highest-scoring two modules, the scores of all other modules fall within one standard deviation of the average score of modules identified on permuted datasets. This confirms that our null models are realistic.

**Figure 2:**
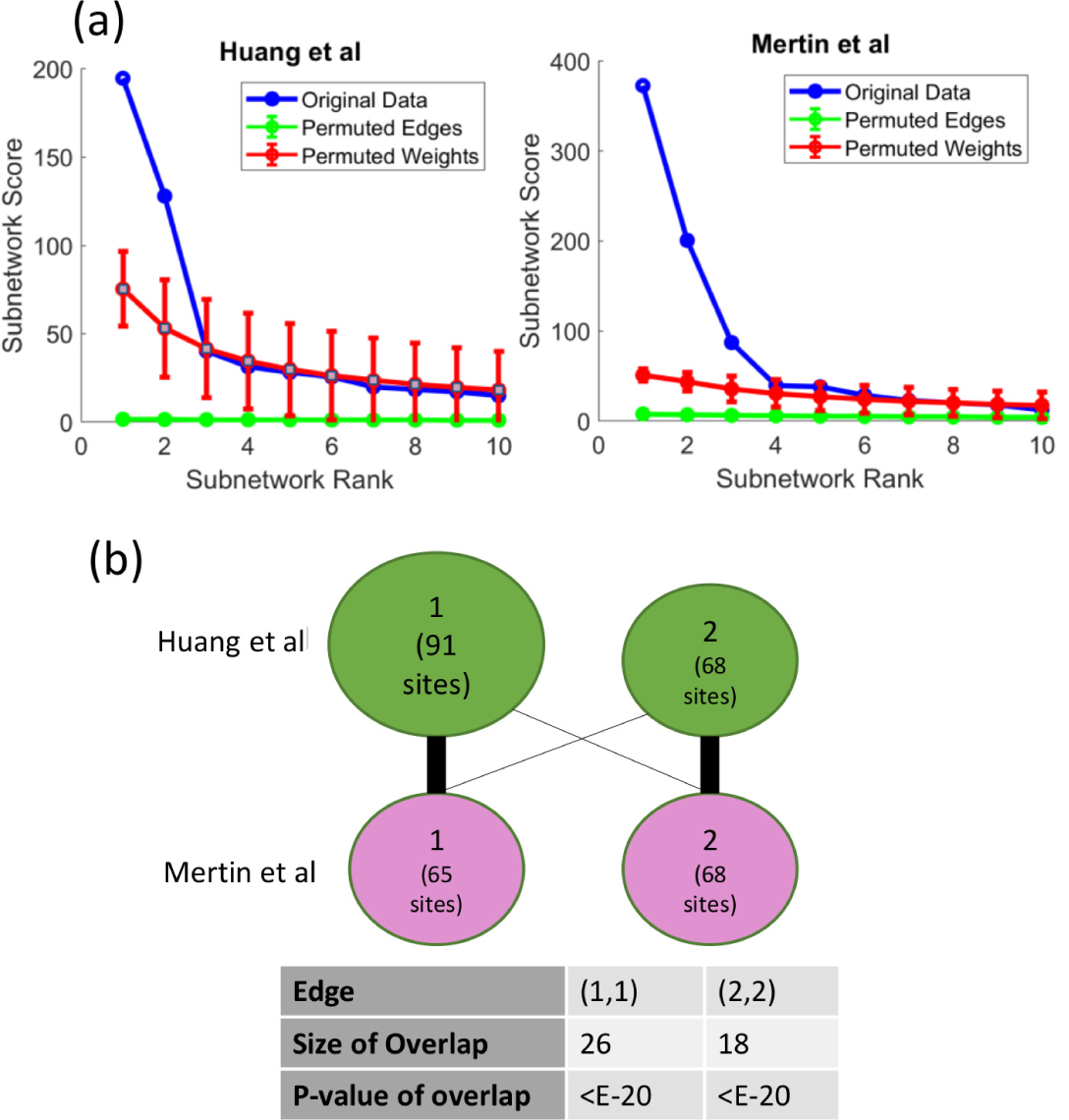
CoPPNet identifies highly significant and reproducible co-phosphorylation (Co-P) modules. (a) Statistical significance of identified sub-networks in two breast cancer datasets. For each dataset, the blue curve shows Co-P scores (y-axis) of the highest scoring 10 sub-networks in decreasing order (rank shown on x-axis). For each rank *i* on the *x-axis*, the red (green) curve and error bar show the distribution of the scores of *i* highest scoring sub-networks in 100 randomized networks obtained by permuting the edge weights (edges). (b) Reproducibility of significant Co-P modules between two independent dataset Huang *et al.* and Mertin *et al.*. The size of the circles indicates the number of phosphosites in each Co-P module, the number in the circle shows its rank among all identified sub-networks. The thickness of the edges represents the significance of the overlap between the two Co-P modules based on hypergeometric test.

We also investigate the reproducibility of the significant modules identified on Huang *et al.* and Mertin *et al.* datasets. In Figure 2(b), the green circles represent the Co-P modules identified on Huang *et al.* dataset and the pink circles represent the Co-P modules identified on Mertin *et al.* dataset. As seen in the figure, there is considerable overlap between the top Co-P modules identified on each dataset; 26 out of the 91 sites in the top Huang *et al.* module and 65 sites in the top Mertin *et al.* module are identical. This overlap is highly statistically significant according to hypergeometric test and is particularly impressive considering that some phosphosites may not be present in a dataset because of the limited coverage of mass spectrometry based phosphoproteomics. Indeed, only 41 of 91 sites in the top Huang *et al.* module are identified in the Mertin *et al.* study, and only 54 of the 65 sites in the top Mertin *et al.* module are identified in the Huang *et al.* study. Many of these phospho-proteins such as *THRAP3 [28]*, NBN [29], *RAD18 [30] and* CDK7 [31] are playing important role in different cancers.

The second top-scoring Co-P modules identified in the two datasets, which are both highly significant (*q <* 0.01), also exhibit significant overlap. Namely, 18 out of the 68 sites in the Huang *et al.* module (of which 33 are present in the Mertin *et al.* dataset) and 68 sites in the Mertin *et al.* module (of which 49 are present in the Huang *et al.* dataset) are identical. Note also that two of the sites in the top Huang *et al.* module are in the second Mertin *et al.* module, and one of the sites in the top Mertin *et al.* module is in the second-ranked Huang *et al.* module. The significant overlap and concordance between the top identified modules across two datasets show that the identified modules are highly reproducible and thus likely to be highly relevant to the dysregulation of signaling processes in breast cancer. We also compare the significant modules with the modules extracted from gene co-expression data published in [32]. The paper reported 11 modules. One of the co-expression modules they reported has 18 genes common with top two significant modules identified by our algorithm.

### Co-P modules identified via unsupervised analysis are associated with breast cancer subtypes

Since the subtype information is not used in the construction of the PSFA network and the assessment of co-phosphorylation, the identification of the Co-P modules is agnostic to the clinically determined subtypes of the samples; i.e. CoPPNet is an *unsupervised* method for the identification of breast-cancer associated signaling modules. However, since the Co-P modules capture co-variation across different samples and this variation can be associated with subtypes, these modules can be informative on subtypes. Motivated by this consideration, we investigate if the phosphorylation levels of phosphosites in the identified modules can differentiate breast cancer subtypes. The results of this analysis for the Huang *et al.* dataset are shown in Figure 3 and S1. Subtype-specific differential phosphorylation of Co-P modules identified on the Mertin *et al.* dataset are presented in Figure S2.

**Figure 3:**
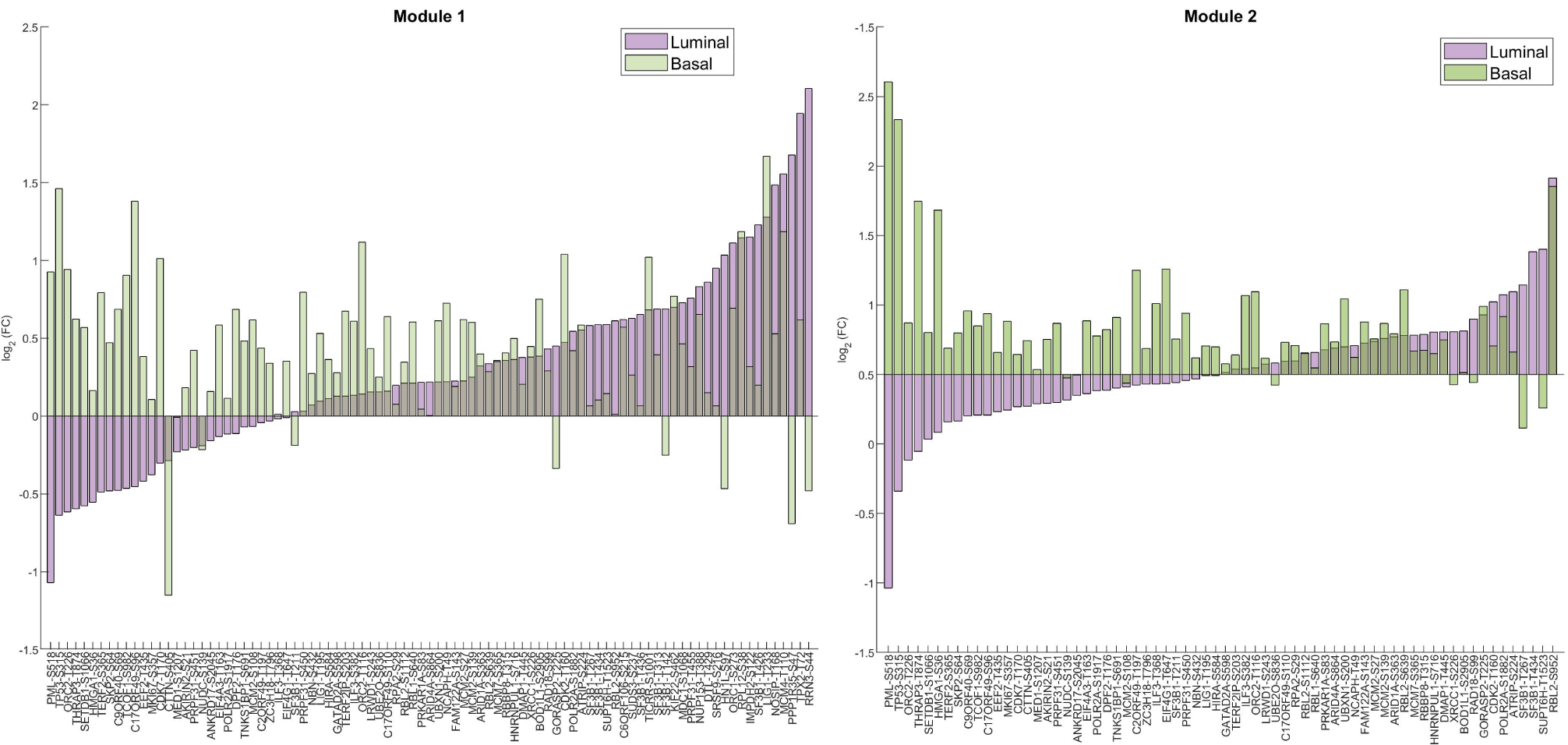
The phosphorylation sites in top Co-P modules identified in Huang et al via unsupervised analysis are associated with breast cancer subtypes. The fold change of the phosphosites in each module are sorted in increasing order of average relative phosphorylation in Luminal samples (purple) with respect to the common reference. The green bars represent the average fold change of phosphorylation in Basal samples.

As seen in Figure 3, top significant Co-P module identified on the Huang *et al.* dataset are highly enriched in phosphosites with significant differential phosphorylation between Luminal and Basal subtypes. There are 14 phosphosites in the top Huang *et al.* module with significant differential phosphorylation between Luminal and Basal subtypes (*p <* 0.05). Eight (*DPF2-T176, THRAP3-T874, TERF2-S365, EIF4A3-T163, SETDB1-S1066, TCOF1-S982, PRPF31-S451, PML-S518*) out of 14 of these sites are hyper-phosphorylated in Basal samples and de-phosphorylated in Luminal samples. For some of the proteins harboring these sites, the differentiation between breast cancer subtypes also has been captured at the level of mRNA expression. For example, *PML* (promyelocytic leukemia) and *SETDB1* (SET Domain Bifurcated 1) are significantly up-regulated in Basal cancers as compared to Luminal cancers, and their expression is related to the survival rate of the patients [33, 34]. We have also compared the relative phosphorylation levels of the sites in the identified modules (Luminal vs. Basal) between the Huang *et al.* and Mertin *et al.* datasets. For module 1, the Pearson, Spearman, and biweight mid- correlation between the relative phosphorylation levels of the sites across the two datasets are respectively 0.37 (*p <* 0.004), 0.37 (*p <* 0.003), and 0.39 (*p <* 0.005). For module 2, the Pearson, Spearman, and biweight mid-correlation between the relative phosphorylation levels of the sites across the two datasets are respectively 0.03 (*p <* 0.41), 0.41 (*p <* 0.01), and 0.25(*p <* 0.01). The result of this analysis is presented in Figure S3.

### Using Co-Phosphorylation Modules for Subtype Prediction

To investigate how the modules can distinguish the subtypes, we use the Co-P modules identified by CoPPNet as features for predicting subtypes on a different dataset. In this analysis, we train a support vector machine (SVM) based classifier for predicting subtypes using the Huang *et al.* dataset as training data. We then test the performance of this classifier on the Mertin *et al.* dataset. For this analysis, for all models that were considered, we restricted the analysis to the sites that were identified in both datasets.

Using this setting, we compared the performance of a model which uses the sites in the significant modules (identified on training data) as features against models that use (i) all the sites that are identified in both datasets (full model), (ii) sites with significant differential phosphorylation levels (p ¡0.05) in the Huang et al. dataset (feature selection using significance of individual sites), and (iii) the top 74 sites according to their differential expression on the Huang et al. dataset (number of features identical to the number of features used by the module-based classifier). The results of this analysis are shown in Table 2. As seen on the table, models that use Co-P modules outperform individual sites and significant sites. While the limited number of samples that are available pose limitations on the generalizability of these results, the improvement provided by Co-P modules demonstrates the promise of Co-P based analysis in differentiating between subtypes.

**Table 2:** Performance of models for subtype prediction using all phosphosites, significant phosphosites(*p*-value ¡ 0.05), and phosphosites in significant Co-P modules.

| All Sites (5476 sites)<br>Accuracy = 46% |  |  | All Significant Sites (621 sites)<br>Accuracy = 46% |  |  |
| --- | --- | --- | --- | --- | --- |
| Real\Predicted | Basal | Luminal | Real\Predicted | Basal | Luminal |
| Basal | 4 | 0 | Basal | 4 | 0 |
| Luminal | 7 | 2 | Luminal | 7 | 2 |

| Significant Sites (74 sites)<br>Accuracy = 46% |  |  | Module 1 & 2 (74 sites)<br>Accuracy = 84% |  |  |
| --- | --- | --- | --- | --- | --- |
| Real\Predicted | Basal | Luminal | Real\Predicted | Basal | Luminal |
| Basal | 4 | 0 | Basal | 4 | 0 |
| Luminal | 7 | 2 | Luminal | 2 | 7 |

### Co-P modules provide a focal point for kinase activity inference

To further understand the contribution of PSFA network and co-phosphorylation analysis, we assess the value added by the Co-P modules to the inference of the differential activity of kinases between Basal and Luminal subtypes. For this purpose, we use the Kinase-Substrate Enrichment Analysis (KSEA) tool, which infers the differential activity of a kinase based on the differential phosphorylation of its substrates [35]. In the kinase enrichment results shown in the Figure 4(a), the analysis is restricted to the target sites of kinases that are in the significant Co-P modules (Basal_*m*_ and Luminal_*m*_) as opposed to all known target sites of the kinase that are identified in the study (Basal_*A*_ and Luminal_*A*_). This analysis infers several kinases with significantly altered activity between the two subtypes. Some of these kinases show different pattern of activity when we limit the KSEA to the phosphosites in the significant modules. To assess the relevance of these kinases, we used the microarray data of breast cancer [26], and ran Kaplan-Meier survival analysis to investigate whether the expression of these kinases is correlated with survival rate. It is well-established that Basal subtype is associated with lower survival rate as compared to Luminal subtype [11]. We observed that, for *AURKA, PRKCI*, higher expression is associated with lower survival rate (Figure 4(b)). KSEA analysis that is restricted to Co-P modules also suggested that these kinases are hyper-active in the Basal samples, however, KSEA on all the phosphosites was not able to capture the association of these kinases with the subtypes. For *MAPK9* and *CDK2*, lower expression is associated with lower survival rate, which is consistent with the kinase activity inferred by restricting to the Co-P modules. The result of this analysis for Mertin *et al.* data is presented in Figure S5.

**Figure 4:**
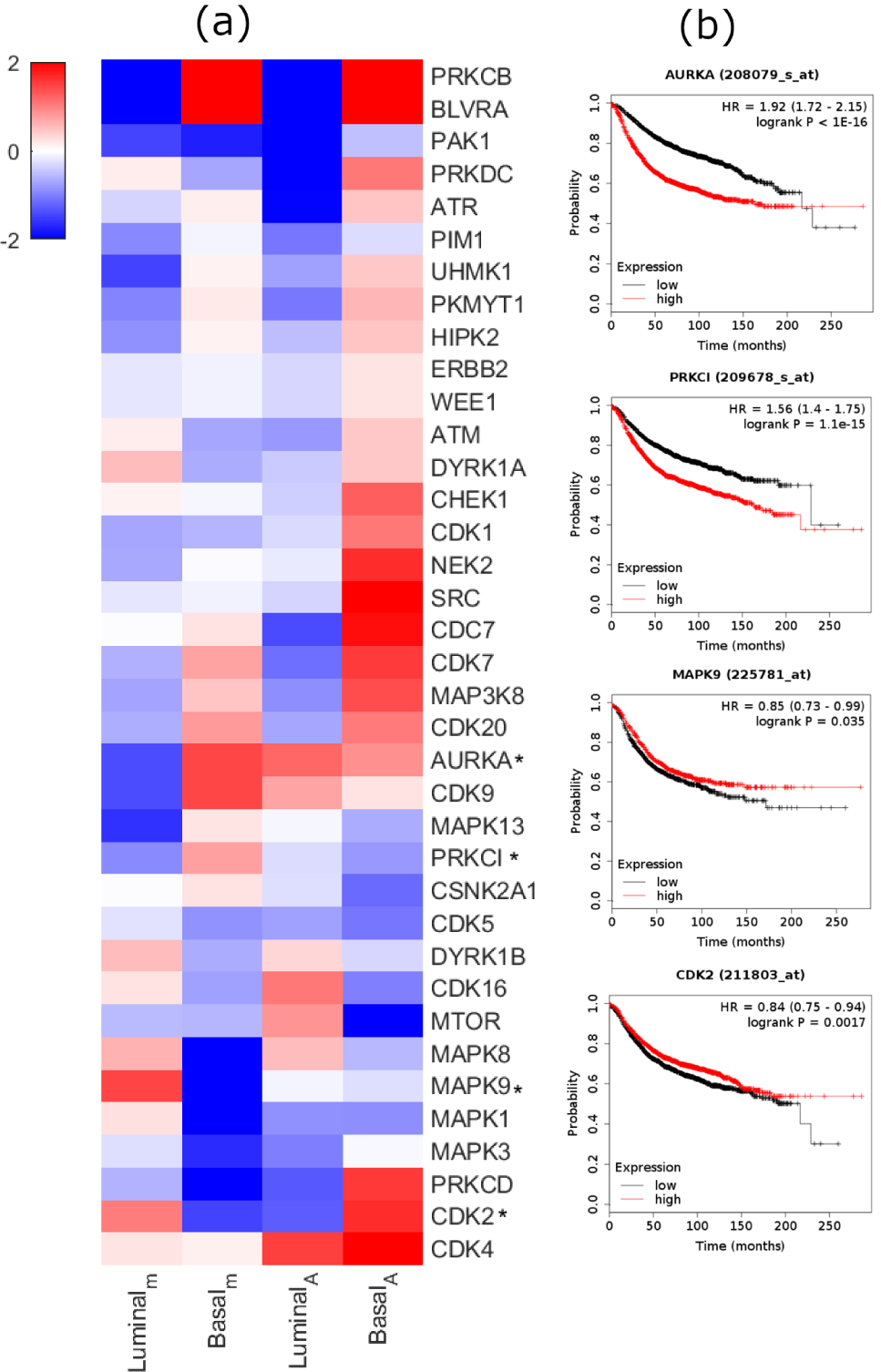
Kinase-substrate enrichment analysis (KSEA) on Co-P modules reveals kinases that are potentially associated with breast cancer subtype and survival. (a) The heatmap compares two different strategies for inferring kinase activity: On the left, the phosphosites utilized to infer kinase activity are restricted to two significant modules identified by CoPPNet(Luminal_*m*_ and Basal_*m*_) on Huang *et al.* dataset. On the right, all phosphosites are used to infer kinase activity (Luminal_*A*_ and Basal_*A*_). The intensity of red indicates the kinases with positive KSEA score (i.e. hyper-active in the respective subtype) and blue indicates the kinases with negative score (i.e. hypo-active in the respective subtype). Kinases that have different patterns of differential activity between subtypes in the modules versus all phosphosites are marked by a star, and their survival analysis using gene expression data is presented in (b).

## 4 CONCLUSION

In this study, we present CoPPNet, a computational method that utilizes large scale phospho-proteomic data for unsupervised identification of phenotype-associated signaling modules in cancer. One important contribution of the proposed method is the construction of the phosphosite functional association (PSFA) network which is a site-centric network that comprehensively incorporates available functional information on phosphorylation sites to enable network-based analysis of phosphorylation data. Our network model treats different types of edges identically. While observation of an edge in different databases would increase the confidence of functional association, we here use the edges only to indicate potential functional association. In future work, it can be useful to investigate the effect of assessing the value of different lines of functional evidence. Our systematic results on two breast cancer datasets show that CoPPNet identifies reproducible subtype-specific signaling modules without requiring knowledge of the sample subtypes. However, this analysis does not account for the tissue-specificity of the phosphorylation data. Overall, this study represents one of the first attempts on utilizing phospho-proteomics to generate reproducible functional readouts of cellular signaling that can be used to characterize the dysregulation of cellular signaling in cancers and development of future therapeutic strategies.

## Acknowledgements

We would like to thank Sean Maxwell, Daniella Schlatzer, and Ming Li for useful discussions.

## Funding

This work was supported in part by US National Institute of Health (NIH) awards R01-LM012980, R01-GM117208 and P30CA043703.

## Notes

https://utrgv-my.sharepoint.com/:b:/g/personal/marzieh_ayati_utrgv_edu/EROMSc_-NslIlnOz-O5UzKIBpsrtzpUNF4K4mB4a8ocyUA?e=jBssAZ

### Competing Interest Statement

The authors have declared no competing interest.

### Summary of Updates

Figure 2 revised; Figure 3 revised; Figure 4 is moved to Supplementary and Figure 4 is updated to new Figure. Table 1 and 2 are added to summarize data.Section "Using Co-Phosphorylation Modules for Subtype Prediction: is added.

## References

[1] Vincentius A Halim, Monica Alvarez-Fernandez, Yan Juan Xu, Melinda Aprelia, Henk WP van den Toorn, Albert JR Heck, Shabaz Mohammed, and Rene H Medema. Comparative phosphoproteomic analysis of check-point recovery identifies new regulators of the dna damage response. Sci. Signal., 6(272):rs9–rs9, 2013.

[2] Rafael Rosell, Teresa Moran, Cristina Queralt, Rut Porta, Felipe Cardenal, Carlos Camps, Margarita Majem, Guillermo Lopez-Vivanco, Dolores Isla, Mariano Provencio, et al. Screening for epidermal growth factor receptor mutations in lung cancer. New England Journal of Medicine, 361(10):958–967, 2009.

[3] James E Butrynski, David R D’adamo, Jason L Hornick, Paola Dal Cin, Cristina R Antonescu, Suresh C Jhanwar, Marc Ladanyi, Marzia Capelletti, Scott J Rodig, Nikhil Ramaiya, et al. Crizotinib in alk-rearranged inflammatory myofibroblastic tumor. New England Journal of Medicine, 363(18):1727–1733, 2010.

[4] Danilo Perrotti and Paolo Neviani. Protein phosphatase 2a: a target for anticancer therapy. The lancet oncology, 14(6):e229–e238, 2013.

[5] S Mohammed, AJ Heck, et al. Phosphoproteomics., 2014.

[6] Tenley C Archer, Tobias Ehrenberger, Filip Mundt, Maxwell P Gold, Karsten Krug, Clarence K Mah, Elizabeth L Mahoney, Colin J Daniel, Alexander LeNail, Divya Ramamoorthy, et al. Proteomics, post-translational modifications, and integrative analyses reveal molecular heterogeneity within medulloblastoma subgroups. Cancer cell, 34(3):396–410, 2018.

[7] Philipp Mertins, DR Mani, Kelly V Ruggles, Michael A Gillette, Karl R Clauser, Pei Wang, Xianlong Wang, Jana W Qiao, Song Cao, Francesca Petralia, et al. Proteogenomics connects somatic mutations to signalling in breast cancer. Nature, 534(7605):55, 2016.

[8] M. Ayati, D Wiredja, D Schlatzer, S. Maxwell, Ming Li, M. Koyuturk, and M.R Chance. Cophosk: A method for comprehensive kinase substrate annotation using co-phosphorylation analysis. PLOS computational biology, 2019.

[9] Yang Yang, Leng Han, Yuan Yuan, Jun Li, Nainan Hei, and Han Liang. Gene co-expression network analysis reveals common system-level properties of prognostic genes across cancer types. Nature communications, 5:3231, 2014.

[10] Jing Liu, Ling Jing, and Xilin Tu. Weighted gene co-expression network analysis identifies specific modules and hub genes related to coronary artery disease. BMC cardiovascular disorders, 16(1):54, 2016.

[11] Saber Fallahpour, Tanya Navaneelan, Prithwish De, and Alessia Borgo. Breast cancer survival by molecular subtype: a population-based analysis of cancer registry data. CMAJ open, 5(3):E734, 2017.

[12] Pablo Minguez, Ivica Letunic, Luca Parca, Luz Garcia-Alonso, Joaquin Dopazo, Jaime Huerta-Cepas, and Peer Bork. Ptmcode v2: a resource for functional associations of post-translational modifications within and between proteins. Nucleic acids research, 43(D1):D494–D502, 2014.

[13] Peter V Hornbeck, Bin Zhang, Beth Murray, Jon M Kornhauser, Vaughan Latham, and Elzbieta Skrzypek. Phosphositeplus, 2014: mutations, ptms and recalibrations. Nucleic acids research, 43(D1):D512–D520, 2014.

[14] Tingting Li, Fei Li, and Xuegong Zhang. Prediction of kinase-specific phosphorylation sites with sequence features by a log-odds ratio approach. Proteins: Structure, Function, and Bioinformatics, 70(2):404–414, 2008.

[15] Andrew Chatr-Aryamontri, Rose Oughtred, Lorrie Boucher, Jennifer Rust, Christie Chang, Nadine K Kolas, Lara O’Donnell, Sara Oster, Chandra Theesfeld, Adnane Sellam, et al. The biogrid interaction database: 2017 update. Nucleic acids research, 45(D1):D369–D379, 2017.

[16] Sara Ballouz, Wim Verleyen, and Jesse Gillis. Guidance for rna-seq co-expression network construction and analysis: safety in numbers. Bioinformatics, 31(13):2123–2130, 2015.

[17] Patrick E Meyer, Frederic Lafitte, and Gianluca Bontempi. minet: Ar/bioconductor package for inferring large transcriptional networks using mutual information. BMC bioinformatics, 9(1):461, 2008.

[18] Lin Song, Peter Langfelder, and Steve Horvath. Comparison of co-expression measures: mutual information, correlation, and model based indices. BMC bioinformatics, 13(1):328, 2012.

[19] Marcus T Dittrich, Gunnar W Klau, Andreas Rosenwald, Thomas Dandekar, and Tobias Müller. Identifying functional modules in protein– protein interaction networks: an integrated exact approach. Bioinformatics, 24(13):i223–i231, 2008.

[20] Marzieh Ayati, Sinan Erten, Mark R Chance, and Mehmet Koyutürk. Mobas: identification of disease-associated protein subnetworks using modularity-based scoring. EURASIP Journal on Bioinformatics and Systems Biology, 2015(1):7, 2015.

[21] Aaron Clauset, Mark EJ Newman, and Cristopher Moore. Finding community structure in very large networks. Physical review E, 70(6):066111, 2004.

[22] Mehmet Koyutürk, Yohan Kim, Umut Topkara, Shankar Subramaniam, Wojciech Szpankowski, and Ananth Grama. Pairwise alignment of protein interaction networks. Journal of computational biology : a journal of computational molecular cell biology, 13:182–99, 04 2006.

[23] Michelle Girvan and Mark EJ Newman. Community structure in social and biological networks. Proceedings of the national academy of sciences, 99(12):7821–7826, 2002.

[24] Bin Zhang and Steve Horvath. A general framework for weighted gene co-expression network analysis. Statistical applications in genetics and molecular biology, 4(1), 2005.

[25] Peter Langfelder and Steve Horvath. Wgcna: an r package for weighted correlation network analysis. BMC bioinformatics, 9(1):559, 2008.

[26] Balazs Györffy, Andras Lanczky, Aron C Eklund, Carsten Denkert, Jan Budczies, Qiyuan Li, and Zoltan Szallasi. An online survival analysis tool to rapidly assess the effect of 22,277 genes on breast cancer prognosis using microarray data of 1,809 patients. Breast cancer research and treatment, 123(3):725–731, 2010.

[27] Kuan-lin Huang, Shunqiang Li, Philipp Mertins, Song Cao, Harsha P Gunawardena, Kelly V Ruggles, DR Mani, Karl R Clauser, Maki Tanioka, Jerry Usary, et al. Proteogenomic integration reveals therapeutic targets in breast cancer xenografts. Nature communications, 8:14864, 2017.

[28] Petra Beli, Natalia Lukashchuk, Sebastian A Wagner, Brian T Weinert, Jesper V Olsen, Linda Baskcomb, Matthias Mann, Stephen P Jackson, and Chunaram Choudhary. Proteomic investigations reveal a role for rna processing factor thrap3 in the dna damage response. Molecular cell, 46(2):212–225, 2012.

[29] Alessandra Di Masi, Francesca Gullotta, Valentina Cappadonna, Loris Leboffe, and Paolo Ascenzi. Cancer predisposing mutations in brct domains. IUBMB life, 63(7):503–512, 2011.

[30] Satoshi Tateishi, Yoshiyuki Sakuraba, Sadaharu Masuyama, Hirokazu Inoue, and Masaru Yamaizumi. Dysfunction of human rad18 results in defective postreplication repair and hypersensitivity to multiple mutagens. Proceedings of the National Academy of Sciences, 97(14):7927–7932, 2000.

[31] Bo Li, Triona Ni Chonghaile, Yue Fan, Stephen F Madden, Rut Klinger, Aisling E O’Connor, Louise Walsh, Gillian O’Hurley, Girish Mallya Udupi, Jesuchristopher Joseph, et al. Therapeutic rationale to target highly expressed cdk7 conferring poor outcomes in triple-negative breast cancer. Cancer research, 77(14):3834–3845, 2017.

[32] Denise M Wolf, Marc E Lenburg, Christina Yau, Aaron Boudreau, and Laura J van‘t Veer. Gene co-expression modules as clinically relevant hallmarks of breast cancer diversity. PloS one, 9(2), 2014.

[33] Arkaitz Carracedo, Dror Weiss, Amy K Leliaert, Manoj Bhasin, Vincent CJ De Boer, Gaelle Laurent, Andrew C Adams, Maria Sundvall, Su Jung Song, Keisuke Ito, et al. A metabolic prosurvival role for pml in breast cancer. The Journal of clinical investigation, 122(9):3088–3100, 2012.

[34] Yuanyuan Jiang, Lanxin Liu, Wenqi Shan, and Zeng-Quan Yang. an integrated genomic analysis of tudor domain–containing proteins identifies phd finger protein 20-like 1 (phf20l1) as a candidate oncogene in breast cancer. Molecular oncology, 10(2):292–302, 2016.

[35] Pedro Casado, Juan-Carlos Rodriguez-Prados, Sabina C Cosulich, Sylvie Guichard, Bart Vanhaesebroeck, Simon Joel, and Pedro R Cutillas. Kinase-substrate enrichment analysis provides insights into the heterogeneity of signaling pathway activation in leukemia cells. Sci. Signal., 6(268):rs6–rs6, 2013.

