## Supplementary File for "Co-Phosphorylation Networks Reveal Subtype-Specific Signaling Modules in Breast Cancer"

### Supplementary Material

**Algorithm 1** Module Identification

```
Input: G = (V, E, w) : PSFA Network;  FC : Fold Change of phosphosites
Output: M : Modules
procedure CoPPNet
    while (All nodes are assigned to a module) do
    //Start the module with the most significant node to make the algorithm deterministic
        m ← max(FC)
        w̄ ← mean(w)
        Score(m) ← 0
        IncompleteModule ← True
        while IncompleteModule do
        //Assess all the neighbors of the nodes in the module and pick the node with the best improvement of score
            N ← All nodes incident to the nodes in m
            NewScore = []
            for all n ∈ N do
                for all u ∈ m do
                    if e_nu ∈ E then
                        s = w_nu − w̄
                    else
                    //Penalize the score if it is not connected to any of the nodes in the module
                        s = 0 − w̄
                    end
                end
                NewScore ← NewScore ⋃ s
            end
            bestNode-Score ← max(NewScore)
            bestNode ← argmax(NewScore)
            if Score(m) + bestNode-Score > Score(m) then
                m ← m ⋃ bestNode
                Score(m) ← Score(m) + bestNode-Score
            else
            //If no additional nodes can improve the module score, remove the nodes from the network and search for the next module
                M ← M ⋃ m
                Remove m from network
                IncompleteModule ← False
            end
        end
    end
```

Table S1: Overlap between different types of edge in the PSFA network.

| Huang et al | FES | KSA | TCK | PPI |
|---|---|---|---|---|
| FES | 7999 | 56 | 58 | 4419 |
| KSA | | 3024 | 462 | 350 |
| TCK | | | 34857 | 438 |
| PPI | | | | 13536 |

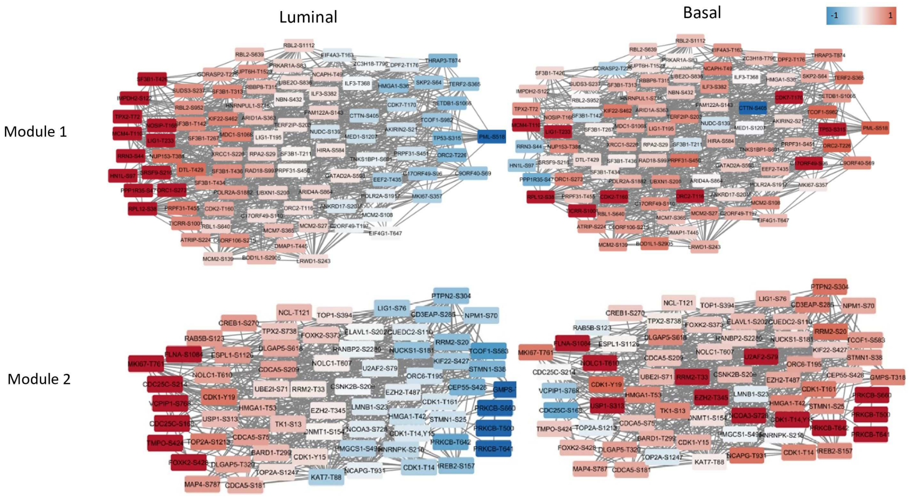


Figure S1: **Top two Co-P module identified in** Huang *et al.***via unsupervised analysis are associated with breast cancer subtypes.** The layout of the module is fixed where the nodes are sorted in decreasing order of average relative phosphorylation in Luminal samples with respect to the common reference. Subtype-specific phosphorylation is shown by node colors. On the left (right) panel, the color of each node indicates the direction of average relative phosphorylation of the phosphosite in Luminal (Basal) samples with respect to the common reference sample, where red indicates hyper-phosphorylation and blue indicates de-phosphorylation. The intensity of the color is identical on the left and on the right, and it indicates the significance of the differential phosphorylation of the site between Luminal and Basal samples.

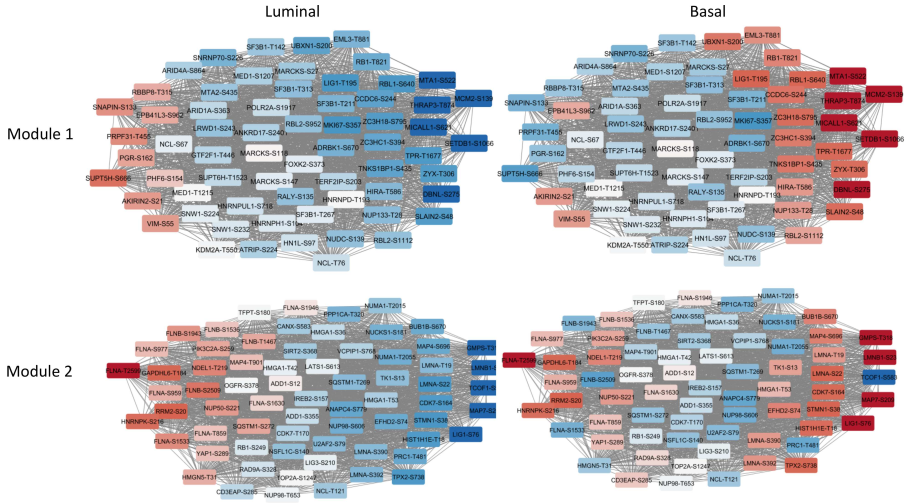


Figure S2: **Co-P modules identified in** Mertin *et al.***via unsupervised analysis are associated with breast cancer subtypes.** (a): Two significant modules identified on Mertin et al are visualized. The layout of each module is fixed where the nodes are sorted in decreasing order of average relative phosphorylation in Luminal samples with respect to the common reference. Subtype-specific phosphorylation is shown by node colors. On the left (right) panel, the color of each node indicates the direction of average relative phosphorylation of the phosphosite in Luminal (Basal) samples with respect to the common reference sample, where red indicates hyper-phosphorylation and blue indicates de-phosphorylation. The intensity of the color is identical on the left and on the right, and it indicates the significance of the differential phosphorylation of the site between Luminal and Basal samples. A node that is colored dark red for Luminal and dark blue for Basal shows a phosphosite that is hyper-phosphorylated in Luminal samples, de-phosphorylated in Basal samples, and exhibits significant differential phosphorylation between the two subtypes. A node that is colored dark red for both Luminal and Basal shows a phosphosite that is hyper-phosphorylated with respect to the reference sample in both subtypes, but still exhibits significant differential phosphorylation between Luminal and Basal samples (the scale is logarithmic with base 2).

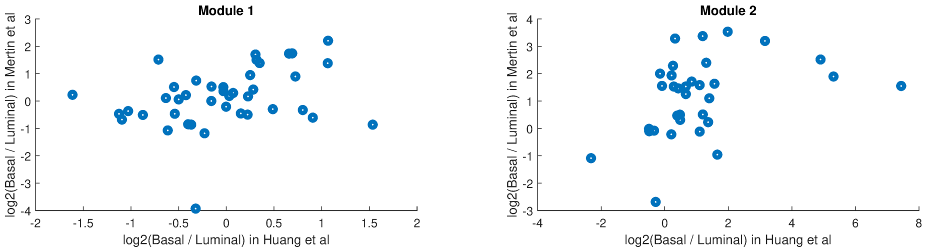


Figure S3: **Relative phosphorylation levels of the sites in the significant Co-P modules (Luminal vs. Basal) between the Huang et al. and Mertin et al. datasets.** For module 1, the Pearson, Spearman, and biweight mid- correlation between the relative phosphorylation levels of the sites across the two datasets are respectively 0.37 ($p < 0.004$), 0.37 ($p < 0.003$), and 0.39 ($p < 0.005$). For module 2, the Pearson, Spearman, and biweight mid-correlation between the relative phosphorylation levels of the sites across the two datasets are respectively 0.03 ($p < 0.41$), 0.41 ($p < 0.01$), and 0.25($p < 0.01$). In these analyses, we use 100 permutation tests by scrambling the relative phosphorylation levels across sites to compute the p-values.

**Effective and direct utilization of phosphorylation data enhances the identification of subtype-associated proteins over protein expression.**

In this section, taking advantage of the availability of mass spectrometry based protein expression data from the samples we use in our experiments, we investigate whether the subtype-specific phosphorylation signatures we identify can be explained by changes in protein expression. For this purpose, we first identify the phosphosites in the top two Co-P modules of the Huang *et al.* dataset with significant differential phosphorylation ($p < 0.05$) between Luminal and Basal subtypes. For each of these sites, we assess the differential expression of the protein harboring the site (if the protein is identified in the protein expression data). The results of this analysis are shown in Figure S4. The result shows while the phosphorylation of these phosphosites is significantly different between two subtypes, most of the the proteins that harbor these sites do not exhibit significant differential protein expression between the two subtypes.

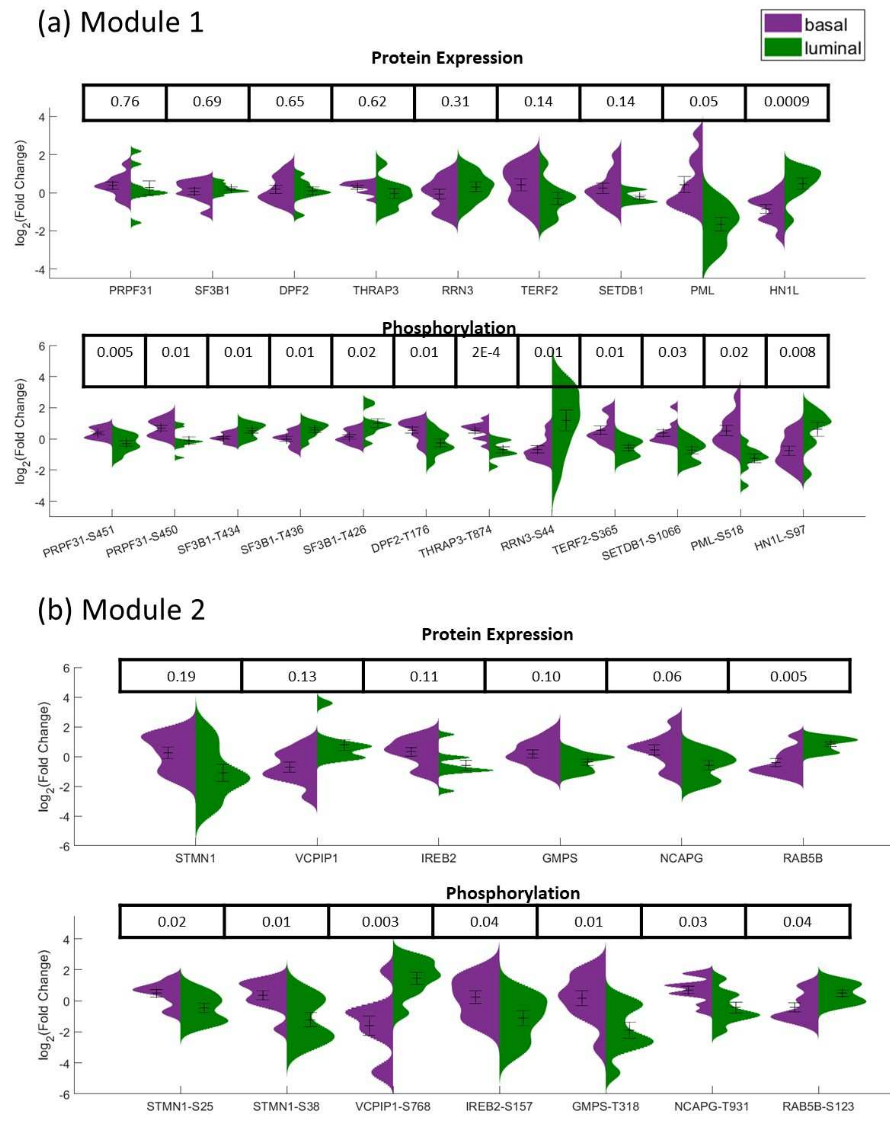


Figure S4: **Effective and direct utilization of phosphorylation data enhances the identification of subtype-associated proteins over protein expression.** For the significant phosphosites of module 1 (a) and module 2 (b) identified on the BC I dataset, the violin plots show the distribution of protein expression and phosphorylation for Luminal (green) versus Basal (purple) samples. The p-value of t-test between these distributions is shown above the plot for each protein and phosphosite.

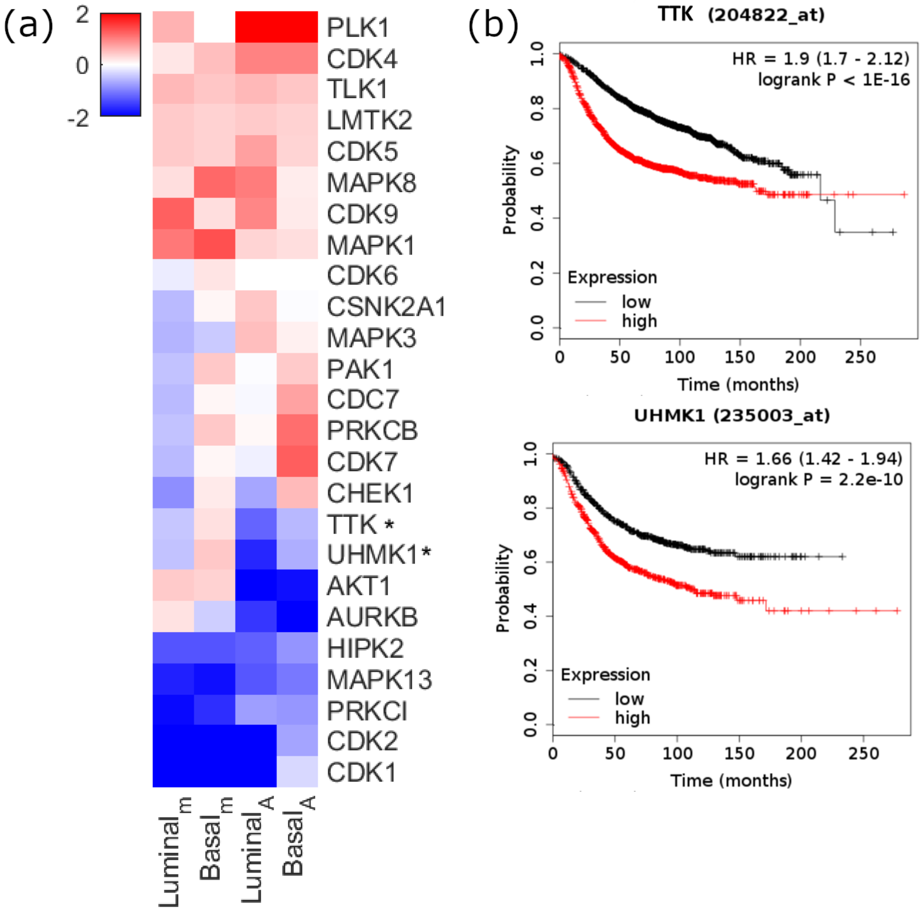


Figure S5: **Kinase-substrate enrichment analysis (KSEA) on Co-P modules reveals kinases that are potentially associated with breast cancer subtype and survival.** (a) The heatmap compares two different strategies for inferring kinase activity: On the left, the phosphosites utilized to infer kinase activity are restricted to two significant modules identified by CoPPNet( Luminal$_m$ and Basal$_m$) on Mertin *et al.* dataset. On the right, all phosphosites are used to infer kinase activity (Luminal$_A$ and Basal$_A$). The intensity of red indicates the kinases with positive KSEA score (i.e. hyper-active in the respective subtype) and blue indicates the kinases with negative score (i.e. hypo-active in the respective subtype). Kinases that have different patterns of differential activity between subtypes in the modules versus all phosphosites are marked by a star, and their survival analysis using gene expression data is presented in (b).

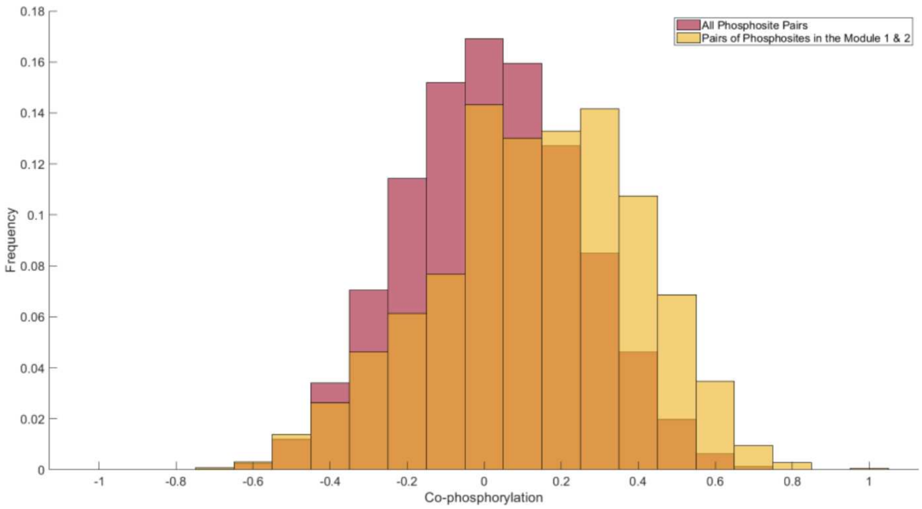


Figure S6: **Co-phosphorylation of the pairs of phosphosites that are within modules against all pairs of phosphosites.** The Kolmogrov-Smirnov test p-value $< E - 20$.

#### Girvan-Newman Result

To apply Girvan-Newman algorithm [1], we considered different thresholds on co-phosphorylation for the inclusion of edges in the network. We used Girvan-Newman modularity score to score the identified modules. Figure S7 shows the distribution of the size of the identified modules and Figure S8 shows the statistical analysis of the identified modules. The overlap of the top identified module using Girvan-Newman algorithm and the top two modules of CoPPNet algorithm is presented in Table . The subnetworks for threshold 0.6-0.9 are presented in Table S3. We have observed that Girvan-Newman algorithm identifies modules that are larger than CoPPNet 's modules and composed of sites on the same protein.

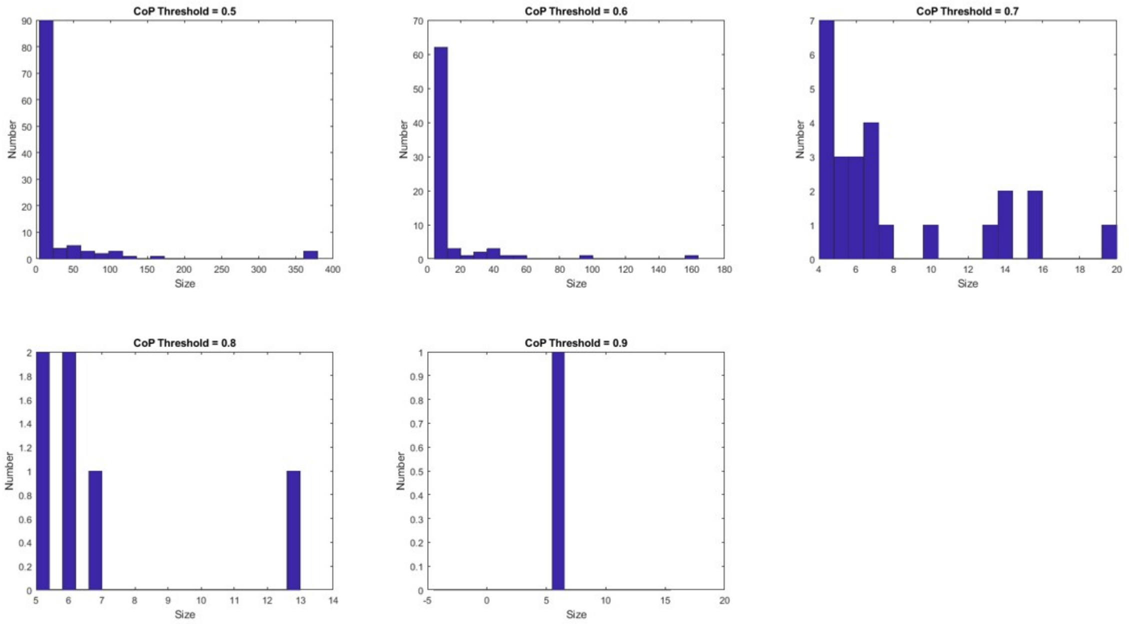


Figure S7: **Distribution of the size of identified modules by Girvan-Newman algorithm for different Co-P threshold.**

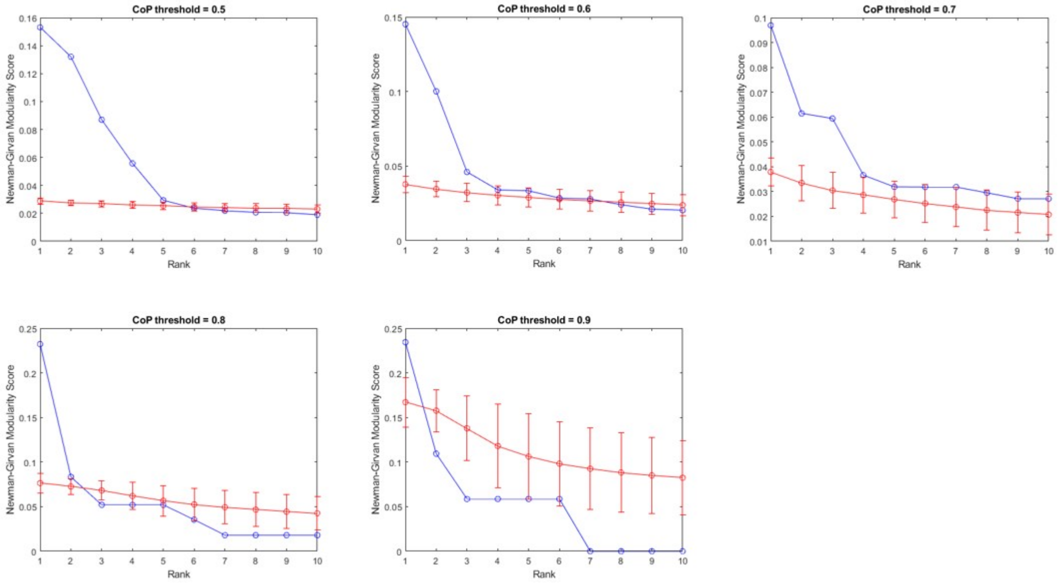


Figure S8: **Statistical significance of identified modules by Girvan-Newman algorithm in** Huang *et al.* **datasets.** For each dataset, the blue curve shows Co-P scores (y-axis) of the highest scoring 10 sub-networks in decreasing order (rank shown on x-axis). Each panel represent different the Co-P threshold (0.5 - 0.9) that is used to include the edge in the network. For each rank $i$ on the *x-axis*, the red curve and error bar show the distribution of the scores of $i$ highest scoring sub-networks in 10 randomized networks obtained by permuting the edge weights.

Table S2: Overlap between the top module of Girvan-Newman algorithm and top two identified modules of CoPPNet algorithm

| threshold | Size of Top Girvan-Newman Module | Module 1 of CoPPNet (91 sites) | Module 2 of CoPPNet (68 sites) |
|---|---|---|---|
| 0.5 | 379 sites | 90 | 59 |
| 0.6 | 164 sites | 42 | 43 |
| 0.7 | 20 sites | 0 | 0 |
| 0.8 | 12 sites | 0 | 0 |
| 0.9 | 6 sites | 0 | 0 |

Table S3: The identified modules by Girvan-Newman algorithm using different Co-P threshold.

| 0.9 | 0.8 | 0.7 | | |
|---|---|---|---|---|
| module 1 | module 1 | module 1 | modue 2 | module 3 |
| 'SFN-S45' | ARHGEF2-S885' | KRT18-S60' | 'ARHGEF2-S885' | 'KAT7-T88' |
| 'YWHAB-S47' | 'CTNNB1-S675' | 'KRT18-S10' | 'CTNNB1-S675' | 'LIG1-S76' |
| 'YWHAE-S46' | 'PAK4-S104' | 'KRT18-S100' | 'GIT2-S415' | 'LIG1-T195' |
| 'YWHAG-S46' | 'PAK4-S148' | 'KRT18-S15' | 'JUP-M89M,S94' | 'MCM2-S108' |
| 'YWHAH-S46' | 'PAK4-S181' | 'KRT18-S23' | 'PAK4-S104' | 'MCM2-S139' |
| 'YWHAQ-S45 | 'PAK4-S267' | 'KRT18-S30' | 'PAK4-S148' | 'MCM2-S27' |
| | 'PAK4-S291' | 'KRT18-S305' | 'PAK4-S181' | 'MCM2-T25,S26' |
| | 'PAK4-S41' | 'KRT18-S319' | 'PAK4-S267' | MCM2-T39' |
| | PAK4-S474' | 'KRT18-S323' | 'PAK4-S267,S291' | MCM3-S756' |
| | 'PAK4-T207' | 'KRT18-S398' | 'PAK4-S291' | MCM3-S756,T758' |
| | 'PXN-S272' | 'KRT8-S281' | 'PAK4-S41' | 'MCM3-T767' |
| | 'RAN-S135' | 'KRT8-S286' | 'PAK4-S474' | 'MCM4-T110' |
| | | 'KRT8-S302' | 'PAK4-T207' | 'MCM6-S13' |
| | | 'KRT8-S358' | 'PKP2-S132,S135' | 'MCM6-S271' |
| | | 'KRT8-S438' | 'PXN-S272' | 'MCM6-S689' |
| | | 'KRT8-S505' | 'RAN-S135' | 'TFDP1-S23' |
| | | KRT8-S62' | | |
| | | KRT8-Y455' | | |
| | | 'PKP2-S151' | | |
| | | 'PKP2-S294' | | |

#### Weighted gene co-expression network analysis (WGCNA) Result

WGCNA uses the topological overlap measure (mutual neighbors) as a proximity measure to hierarchically cluster genes into network modules [2, 3]. We used power of 3 for soft thresholding. WGCNA does not score and rank the identified subnetworks, therefore, we do not assess the statistical significance of the identified subnetworks. Figure S9 presents the distribution of the size of identified modules. As seen in the figure, some of the identified modules are larger than the modules identified by CoPPNet. Four identified modules are presented in Table S4. As seen in the table, WGCNA tends to cluster the phosphosites on the same protein together. The maximum overlap between the significant modules identified using CoPPNet and WGCNA are 21 and 37 phosphosites for the first and second module, respectively.

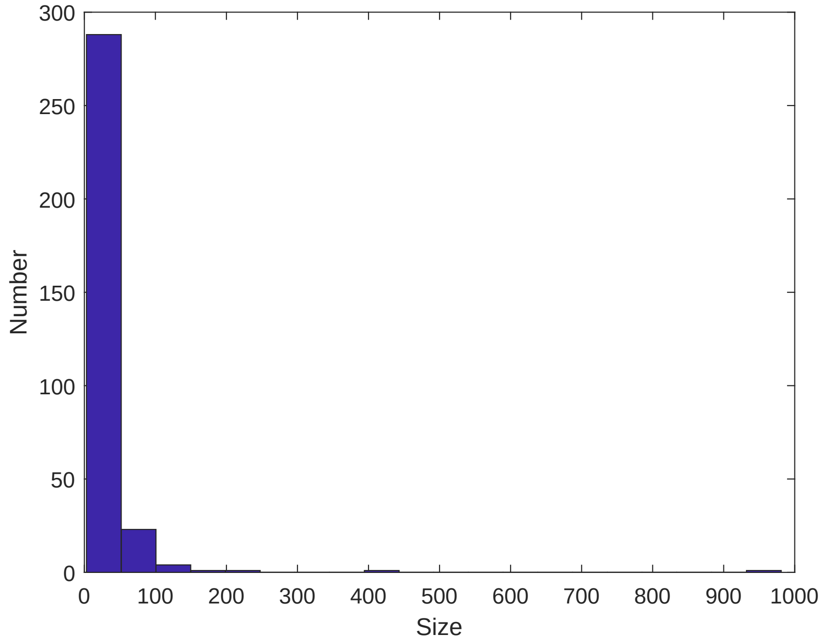


Figure S9: **Distrubtion of size of the modules identified by WGCNA algorithm in** Huang *et al.* **data**

Table S4: Four randomly selected modules from WGCNA.

| Module 1 | Module 2 | Module 3 | Module 5 |
|---|---|---|---|
| 'MAP3K2-S153' | 'AGAP2-S638' | 'MED1-S1156' | 'CTTN-S11' |
| 'MAP3K2-S164' | 'AGAP2-S808' | 'MED1-S1192' | 'CTTN-S261' |
| 'MAP3K2-S239' | 'ITPR1-S1752' | 'MED1-S1207,T1215' | 'CTTN-S298' |
| 'MAP3K2-S331,S344' | 'MYH9-T725' | 'MED1-S1401' | 'CTTN-S447' |
| 'MAP3K2-S344' | 'NUMA1-S1225' | 'MED1-S1433' | 'CTTN-S47' |
| 'MAP3K2-S347' | 'NUMA1-S169' | 'MED1-S1481,S1482' | 'CTTN-T323' |
| 'MAP3K2-S514' | 'NUMA1-S1724' | 'MED1-S664' | 'CTTN-T401' |
| 'ZAK-S275' | 'NUMA1-S1757' | 'MED1-T1051' | 'CTTN-T401,S405' |
| 'ZAK-S593' | 'NUMA1-S1769' | 'MED1-T805' | |
| 'ZAK-S637' | 'NUMA1-S1789' | 'MED24-S862' | |
| 'ZAK-S648' | 'NUMA1-S1800' | 'MED24-S873' | |
| 'ZAK-S727' | 'NUMA1-S1833,S1834' | 'ATM-T1884' | |
| 'ZAK-T628' | 'NUMA1-S1862' | 'CHEK2-T426' | |
| | 'NUMA1-S1945' | 'H2AFX-S140' | |
| | 'NUMA1-S1969' | 'AKAP1-S169' | |
| | 'NUMA1-S1991' | 'AKAP1-S429' | |
| | 'NUMA1-S203' | 'AKAP1-S445' | |
| | 'NUMA1-S271' | 'AKAP1-S592' | |
| | 'NUMA1-S395' | 'AKAP1-T533' | |
| | 'NUMA1-S820' | 'AKAP1-T70' | |
| | 'NUMA1-T2000' | 'ARFGEF1-S1079' | |

### References

[1] Michelle Girvan and Mark E Newman  Community structure in social and biological networks. *Proceedings of the national academy of sciences*, 99(12), 2002.

[2] Bin Zhang and Steve Horvath. A general framework for weighted gene co-expression network analysis. *Statistical applications in genetics and molecular biology*, 4(1), 2005.

[3] Peter Langfelder and Steve Horvath. WGCNA: an R package for weighted correlation network analysis. *BMC bioinformatics*, 9(1), 2008
